# Negative Charges, Not Necessary Phosphorylation, are Required for Ligand Recognition by 14-3-3 Proteins

**DOI:** 10.1101/2024.09.16.613320

**Authors:** Seraphine Kamayirese, Laura A. Hansen, Sándor Lovas

## Abstract

Protein-protein interactions involving 14-3-3 proteins regulate various cellular activities in normal and pathological conditions. These interactions have mostly been reported to be phosphorylation-dependent, but the 14-3-3 proteins also interact with unphosphorylated proteins. In this work, we investigated whether phosphorylation is required, or, alternatively, whether negative charges are sufficient for 14-3-3ε binding. We substituted the pThr residue of pT(502-510) peptide by residues with varying number of negative charges, and investigated binding of the peptides to 14-3-3ε using MD simulations and biophysical methods. We demonstrated that at least one negative charge is required for the peptides to bind 14-3-3ε while phosphorylation is not necessary, and that two negative charges are preferable for high affinity binding.

## Introduction

Protein phosphorylation, one of the most common post-translational modifications, plays an important role in the regulation of cellular functions. Protein kinases commonly phosphorylate seryl, threonyl and tyrosyl residues of proteins, and this modification not only affects structural properties of the proteins but also their functions [1–3]. Phosphorylation modulates protein-protein interactions, and some proteins recognize binding motifs containing phosphorylated amino acid residues [4–8].

14-3-3 proteins are a family of adapter proteins that are ubiquitously expressed in eukaryotic cells. The 14-3-3 proteins have many binding partners with a broad range of functions. Thus, these proteins are involved in various cellular activities in both physiological and pathological conditions [9–12]. The 14-3-3ε isoform is upregulated and linked to abnormal cell growth in renal cancer [13]. Reduced expression of 14-3-3ε has been reported in gastric cancer [14]. The 14-3-3 proteins have also been associated with progression of larynx squamous cell carcinoma (LSCC) [15], and small cell lung cancer [16]. The 14-3-3 isoforms, ε, ζ and γ are expressed and mislocalized to cytoplasm in cutaneous squamous cell carcinoma (cSCC), where a heterodimer of the ε with either ζ or γ interacts with the cell cycle regulator, cell division cycle 25 A (CDC25A) to suppress apoptosis [17–19].

It is widely reported that interactions between 14-3-3 proteins and their binding partners are through binding motifs comprising a phosphoseryl (pSer) or a phosphothreonyl (pThr) amino acid residues [20,21]. The 14-3-3 proteins recognize two binding motifs, RSXp-SXP and RXY/FXpSXP; pS is pSer, and X is any amino acid [20]. Apoptosis signal-regulating kinase 1 (ASK1) [22], Raf-1 [23,24], CDC25A [17], cytosine-anenosine-adenosine-thymidine (CCAAT)-enhancer binding protein [25], and cystic fibrosis transmembrane conductance reg-ulator [8] are among the phosphorylated binding partners of 14-3-3 proteins. However, not all 14-3-3 interactions with their binding partners are phosphorylation-dependent, as they also bind proteins such as CDC25B [26] and exoenzyme S [27,28] in a phosphorylation-independent manner.

Over the years, the binding mechanism of various phosphopeptides to 14-3-3 proteins has been studied using experimental and computational methods [20,29–31]. Muslin and colleagues [24] showed the phosphorylation-dependent binding of the Raf-1 derived peptide (pS-Raf-259) to 14-3-3ζ. The peptide occupies the amphipathic binding groove of 14-3-3ζ, and residues of the protein interacting with the peptide were identified [32], similar interaction were identified between 14-3-3ζ and phosphorylated myeloid leukemia factor 1 peptide [33]. Furthermore, the human SOS1-derived peptide binds 14-3-3ζ in a similar manner as pS-Raf-259 does [34]. In our previous work [19,30,31], we also developed CDC25A derived phosphopeptides that bind 14-3-3ε, and we identified basic and aromatic amino acid residues (Lys^50^, Arg^57^, Arg^130^, Tyr^131^) in the binding pocket of 14-3-3ε (Figure 1) that interact with phosphorylated amino acid residues.

**Figure 1.**
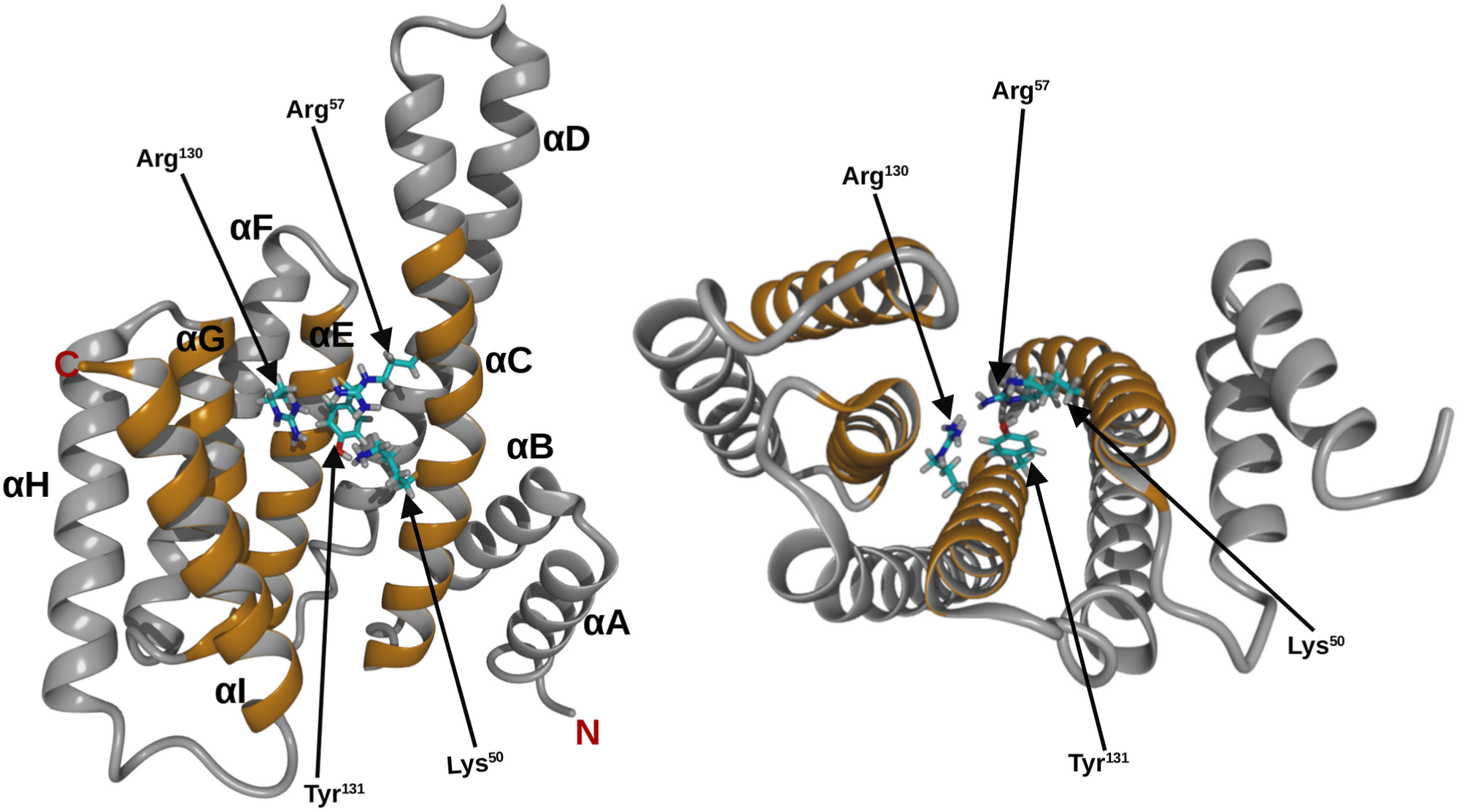
Ribbon structure of 14-3-3ε monomer. Top view (left), side view (right), helices are labeled A to I, from *N-* to *C-* terminal. Helices that form the binding pocked are shown in orange. Amino acid residues of 14-3-3ε that interact with the phospharylated residue of the ligand are shown as stick.

Phosphorylation-independent interactions between 14-3-3 proteins and peptides have also been elucidated. R18, the unphosphorylated peptide that has two negatively charged amino acid residues, discovered by phage display designed for 14-3-3τ binding has been shown to interact with 14-3-3 proteins in a manner reminiscent of phosphorylated peptides. Although, R18 has additional hydrophobic interactions with the protein [23,32]. Ottmann and colleagues [27] demonstrated that the exoenzyme S derived peptide interacts with 14-3-3 protein through ionic and hydrophobic interactions.

We have developed phosphopeptides that inhibit interactions between CDC25A and 14-3-3ε, and induce apoptosis of cSCC in animals [19]. We further improved binding affinities of the peptides for 14-3-3ε by 6.5 fold [30]. Although these phosphopeptides have high affinities for 14-3-3ε, it is likely that they are susceptible to dephosphorylation as various phosphatases have been shown to dephospharylate phosphopeptides in solution [35–37]. Therefore, we proposed replacing the pThr residue of the parent peptide (pT(502-510)) with pThr mimics to potentially improve their resistance to phosphatases. It has long been known that 14-3-3 protein interactions with their binding partners are dependent on phosphorylation of the binding partner. Thus, we aimed to elucidate whether the binding of peptides is dependent on the presence of negatively charged amino acid residues, and not necessarily their phosphorylation.

Here, we studied binding between 14-3-3ε and peptide analogs in which the pThr residue in our previously studied peptide, pT(502-510) [30], was substituted with residues carrying varying number of negative charges. We used biophysical methods to show that at least one negative charge is required for peptide binding to 14-3-3ε.

## Materials and Methods

### Molecular Dynamics (MD) simulations

#### Peptide – 14-3-3ε complexes preparation

The starting structure was obtained from our previously studied 14-3-3ε – pT(502-510) complex [30]. To obtain the various peptide analogs, pThr^507^ amino acid residue in the pT(502-510) was substituted with Thr, Glu, Gla, sThr or Pmb amino acid residues (Figure 2). Non-standard residues were built in the YASARA program [38]. Binding of the peptides to 14-3-3ε was studied using MD simulations.

**Figure 2.**
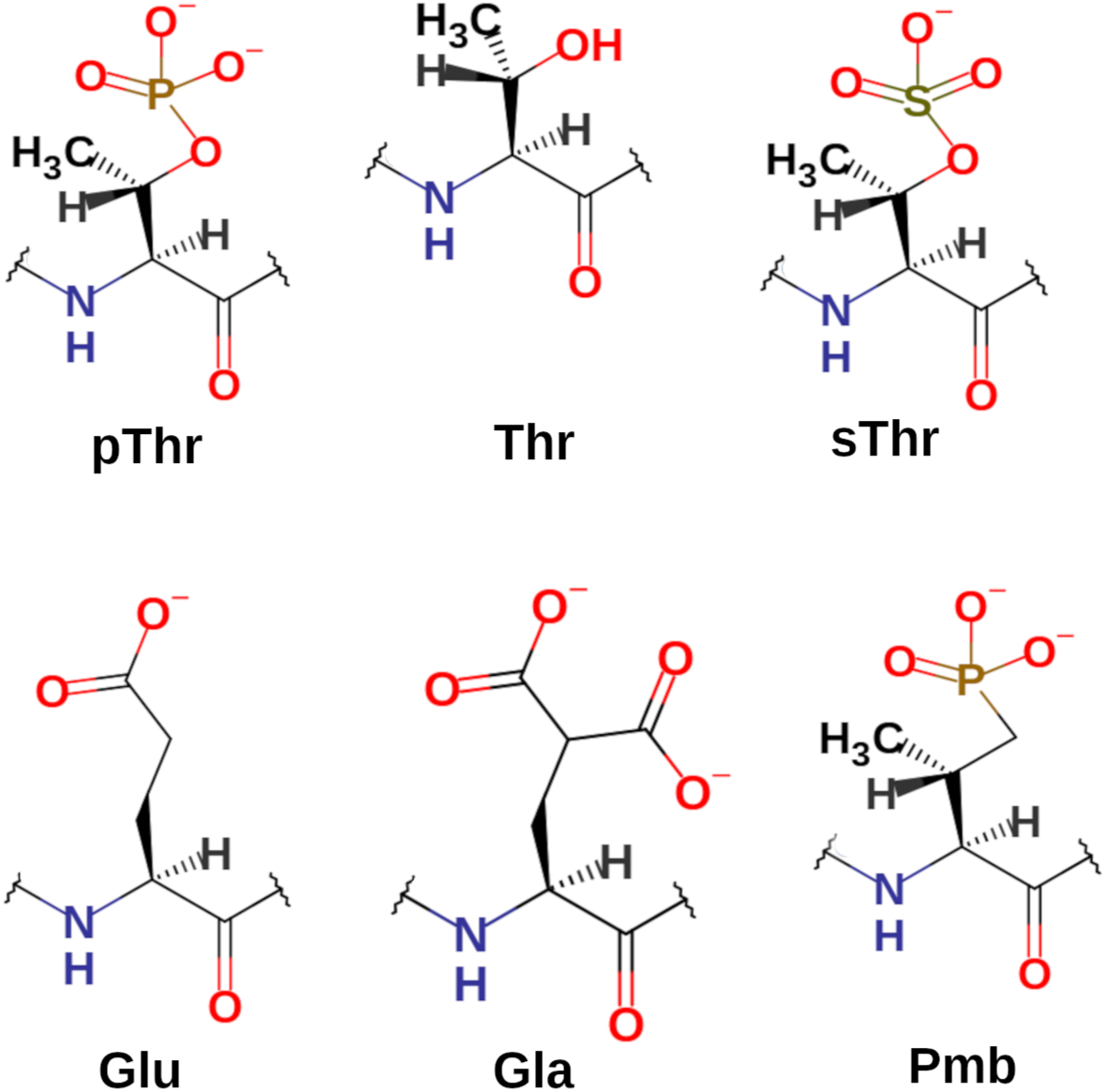
Structure of amino acid residues. Amino acid residues in position 507 of the peptide analogs have varying negative charge number from 0 to −2. pThr, Phosphothreonyl; Thr, threonyl; sThr, sulfothreonyl; Glu, glutamic acid residue; Gla, γ-carboxyglutamic acid residue; Pmb, 2-Amino-3-methyl-4-phosphonobutanoic acid residue.

#### MD simulations of peptide – 14-3-3ε complexes

MD simulations of the peptide – protein complexes were performed using the AMBER-FB15 force field as implemented in YASARA [39]. Initially, the 14-3-3ε – peptide complex was solvated with water in a cubic simulation box with minimal distance of 1.2 nm between the edge of the box and the complex. Then the system was energy minimized using standard parameters in YASARA. Then the energy-minimized structures of 14-3-3ε-peptide complexes were solvated again with water molecules and 150 mM NaCl, and the structure of the system was simulated using *md_run* macro of YASARA. The 500 ns simulations were performed at 310 K and 1 atm pressure, and the trajectories were saved in xtc format that is compatible with GROMACS [40]. The simulations were performed in two steps, (1) a 1 ns simulation using YASARA macro in the YASARA graphical user interface, (2) then the simulation was continued in a computer background to generate a 499 ns trajectory using *md_run* macro of YASARA.

### Trajectory analysis

Using YASARA, the first frames of the trajectories were converted to PDB format and used as topology files for analysis using analysis utilities of GROMACS 2022.5 [40]. The trajectories were processed using *trjconv* module of GROMACS. The *rms* module of GROMACS was used to calculate Cα atoms root-mean-square deviation (RMSD). VMD software [41] was used to calculate salt-bridges and H-bonds between peptides and 14-3-3ε; 0.8 nm was used as the oxygen–nitrogen cut-off distance for salt bridges, and the donor–acceptor distance was set to 0.4 nm for H-bonds. The *sasa* module of GROMACS was used to calculate the solvent accessible surface area (SASA). The interfacial surface area (ISA) [42] was calculated as follow:

ISA = (SASA_14-3-3ε_ + SASA_peptide_) – SASA_14-3-3ε-peptide complex_. The van der Waals (VDW) surface area of residues were calculated using YASARA.

#### Peptides

The [sThr^507^]pT(502-510) peptide was from Biosynth International Inc.(Louisville KY, USA), other peptide analogs were synthesized in house. All *N*-α-Fmoc-protected amino acids were from CEM Corporation (Matthews NC, USA), except, *N*-alpha-(9-Fluorenylmethyloxycarbonyl)-gamma-Carboxy-L-glutamic-acid-gamma-di-t-butyl ester, (Fmoc-L-Gla(OtBu)2-OH) and (2S,3R)-2-((9-fluorenylmethyloxycarbonyl)amino)-4-(di-t-butylphosphonomethyl)-3-methylbutyric acid, (Fmoc-L-Pmb(tBu)2-OH) that were from Iris Biotech (Adalbert-Zoellner-Str 1 D-95615 Marktredwitz, Germany). Dimethylformamide (DMF) and *N*,*N′*-Diisopropylcarbodiimide (DIC) were from Sigma-Aldrich, the Rink amide protide resin and Oxyma Pure were from CEM.

### Peptide synthesis

The peptides were synthesized using CEM Liberty blue 2.0 microwave peptide synthesizer, in a 0.1 mmol scale on Rink amide protide resin using standard Fmoc chemistry with Boc, tBut, Bzl, and Pbf side chain protections. A 1 min Fmoc deprotection was carried out using 20% piperidine in DMF, and 2 min coupling was performed in DIC/Oxyma Pure at 90 °C, except for the (Fmoc-L-Pmb(tBu)2-OH) that was coupled for 4 min. Peptide cleavage from the resin and side chains deprotection were simultaneously performed by stirring the peptide-resin in a cocktail containing TFA/Thioanisol/phenol/TIS/H2O/DODT in 81.5:5:5:1:5:2.5 (v/v/v/v/v/v) for 30 min on ice and then 3.5 hours at room temperature. The resin was then separated from peptide by filtration and the peptide was precipitated using ice-cold ether. Crude peptides were purified by reverse-phase HPLC using C8 column (Phenomenex, Aeris 100 Å, 5 μM, 250 x 10 mm). A 0.1% TFA aqueous solution (v/v solvent A) and a 0.09% TFA acetonitrile (solvent B) were used to elute the peptides with a linear gradient of solvent B of 3 to 60% over 60 min at a flow rate of 4 mL/min. Identities of the peptides were confirmed by Thermo Q-Exactive Orbitrap mass spectrometer.

### Differential scanning fluorimetry (DSF)

The (His)6-14-3-3ε was from Novus Biologicals (Centennial, CO, USA). Thermal unfolding of 14-3-3 protein and 14-3-3 – peptide complexes were performed as previously described [30]. Briefly, thermal unfolding was monitored using SYPRO-orange dye (Invitrogen, Carlsbad, CA, USA) in BioRad CFX384 Touch real-time PCR instrument (Hercules CA, USA). The protein and protein-peptide complexes were dissolved in TRIS buffer (150 mM NaCl, 20 mM TRIS, 1 mM DTT, pH 7.4) to the final concentrations of 30 μM of peptide, 2 μM of 14-3-3ε, and 1000x dilution from SYPRO-Orange dye stock from the manufacturer. Melting temperature (T_m_) was determined from the first derivative of unfolding curve and change in melting temperature (ΔT_m_) was calculated as the difference between T_m_ of protein-peptide complex and Tm of protein alone. At least 3 replicates were performed per peptide.

### Surface plasmon resonance (SPR)

To obtain binding affinity (K_D_) of peptides for 14-3-3ε, SPR was performed as before [30]. Briefly, recombinant His6-14-3-3ε was immobilized on activated Biacore NTA chip (Cytiva, Marlborough MA, USA), at a flow rate of 5 μL / min for 10 min and then washed with the running buffer (20 mM Tris, 150 mM NaCl, 50 μM EDTA, 0.005% Tween 20, pH 7.4) for 20 min. Following protein immobilization, solutions of peptides were injected at increasing concentrations ranging from 4 to 500 uM (3.9,7.8, 15.6, 31.2, 62.5, 125, 250 and 500 μM) for all analogs, except [Pmb^507^]pT(502-510) where concentrations ranging between 0.002 μM and 0.25 μM (0.002, 0.004, 0.008, 0.016, 0.031, 0.063, 0.125 and 0.25 μM) were used. The injections were carried out at a flow rate of 30 μL/min for 1 min, and dissociation time was 3 min in the running buffer. Steady state affinities of the peptide were obtained by fitting the data using Biacore Insight Evaluation Software (version 5.0.18.22102). For each peptide, at least 5 replicates were done.

## Results

In the present work, we elucidated the binding of peptide analogs of our previous studied peptide pT(502-510) [30], in which the pThr amino acid residue in position 507 was substituted with amino acid residues with uncharged Thr, the singly charged sulfothreonine (sThr) and Glu, or with the doubly charged γ-carboxyl glutamic acid (Gla) and 2-Amino-3-methyl-4-phosphonobutanoic acid (Pmb) residues (Table 1). Their chemical structures are shown in Figure 2. The sThr is a phosphatase resistant pThr mimetic with one negative charge, and Pmb is a phosphatase resistant pThr mimetic with two negative charges. For comparison, pThr residue, in our published work, [30] is also shown. Binding of each peptide analog to 14-3-3ε was computationally investigated in 500 ns MD simulations.

**Table 1.** Binding affinities of the peptides for 14-3-3ε, and interacting surface area (ISA) between 14-3-3ε and the peptides. Residues that substituted pThr of the parent peptide (pT(502-510)) are highlighted in red. Ac, and NH_2_, acetyl and amide, respectively, protecting groups. K_D_ values are average ± SD of n ≥ 5.

| Peptide | Amino acid sequence | K<sub>D</sub> / μM | ISA / nm<sup>2</sup> |
| --- | --- | --- | --- |
| pT(502-510)* | Ac-Arg-Thr-Lys-Ser-Arg-pThr-Trp-Ala-Gly-NH<sub>2</sub> | 0.045 ± 0.024* | 15.47 |
| [Thr<sup>507</sup>]pT(502-510) | Ac-Arg-Thr-Lys-Ser-Arg-Thr-Trp-Ala-Gly-NH<sub>2</sub> | No binding | 14.12 |
| [sThr<sup>507</sup>]pT(502-510) | Ac-Arg-Thr-Lys-Ser-Arg-sThr-Trp-Ala-Gly-NH<sub>2</sub> | 86.71 ± 8.07 | 13.67 |
| [Glu<sup>507</sup>]pT(502-510) | Ac-Arg-Thr-Lys-Ser-Arg-Glu-Trp-Ala-Gly-NH<sub>2</sub> | 141.75 ± 24.96 | 16.23 |
| [Gla<sup>507</sup>]pT(502-510) | Ac-Arg-Thr-Lys-Ser-Arg-Gla-Trp-Ala-Gly-NH<sub>2</sub> | 24.06 ± 8.43 | 15.52 |
| [Pmb<sup>507</sup>]pT(502-510) | Ac-Arg-Thr-Lys-Ser-Arg-Pmb-Trp-Ala-Gly-NH<sub>2</sub> | 0.009 ± 0.001 | 17.27 |

\*From our previous work [30]

The Cα-atoms RMSD indicated that the peptides did not dissociate from 14-3-3ε – peptide complexes throughout simulations (Figure S1). All 14-3-3ε – peptide complexes under-went structural rearrangement with RMSD between 0.15 nm and 0.4 nm, except the 14-3-3ε – [Gla^507^]pT(502-510) complex that had RMSD of 0.15–0.6 nm (Figure S1). In all complexes, peptides showed less fluctuation than 14-3-3ε, except the [Pmb^507^]pT(502-510) peptide that showed higher fluctuation between 250 ns and 450 ns. Analysis of the interfacial surface area (ISA) between 14-3-3ε and peptide showed that the 14-3-3ε – peptide complexes had ISA of 13.7–17.3 nm^2^ (Table 1). The [Pmb^507^]pT(502-510) containing complex showed the largest ISA, while the [sThr^507^](502-510) containing complex had the smallest ISA. This suggests that all the peptides had a degree of interaction with 14-3-3ε during simulations. We further determined VDW surface area of the residue in position 507 of each peptide in each 14-3-3ε – peptide complexes. The results indicated that VDW surface area of the residues was between 1.386 nm^2^ and 1.954 nm^2^ (Figure S2). The isosteric replacements, Pmb, Gla, and sThr residues had VDW surface area of 1.954 nm^2^, 1.824 nm^2^ and 1.816 nm^2^, respectively, and the nonisosteric replacements, Glu and Thr had VDW surface area of 1.605 nm^2^ and 1.386 nm^2^, respectively.

We identified several residues of the 14-3-3ε-peptide complex that interact with residues introduced in position 507 (Figure 3, and Table S1). In the 14-3-3ε – [Thr^507^]pT(502-510) complex, no interactions involving Thr^507^ residue were observed. However, residues Arg^502^, Lys^504^, Ser^505^, Arg^506^ and Trp^508^ of the peptide formed H-bonds with residues Lys^123^, Glu^134^, Glu^183^, Asn^176^, Asp^226^ and Asn^227^ of 14-3-3ε (Table S1). In all other 14-3-3ε – peptide complexes, H-bonds were observed between the 507 residue of the peptides and Lys^50^, Arg^57^, Arg^130^ and Try^131^ of 14-3-3ε (Table S1). Pmb residue in the [Pmb^507^]pT(502-510) peptide formed the longest lasting H-bonds with these residues of 14-3-3ε, followed by Gla residue in [Gla^507^]pT(502-510). Additionally, Pmb formed H-bonds with Asn^176^, Leu^175^, Val^179^ and Leu^223^ (Figure 3, and Table S1). The singly charged sThr and Glu in [sThr^507^]pT(502-510) and [Glu^507^]pT(502-510), respectively, showed less lasting H-bonds with Lys^50^, Arg^57^, Arg^130^ and Try^131^ residues of 14-3-3ε, than Gla and Pmb. Analysis of ionic interactions revealed no interactions between the 507 residues of the peptides with residues of the 14-3-3ε, except the Glu residue in [Glu^507^]pT(502-510) peptide that formed salt-bridges with Lys^50^, Lys^123^, Arg^130^.

**Figure 3.**
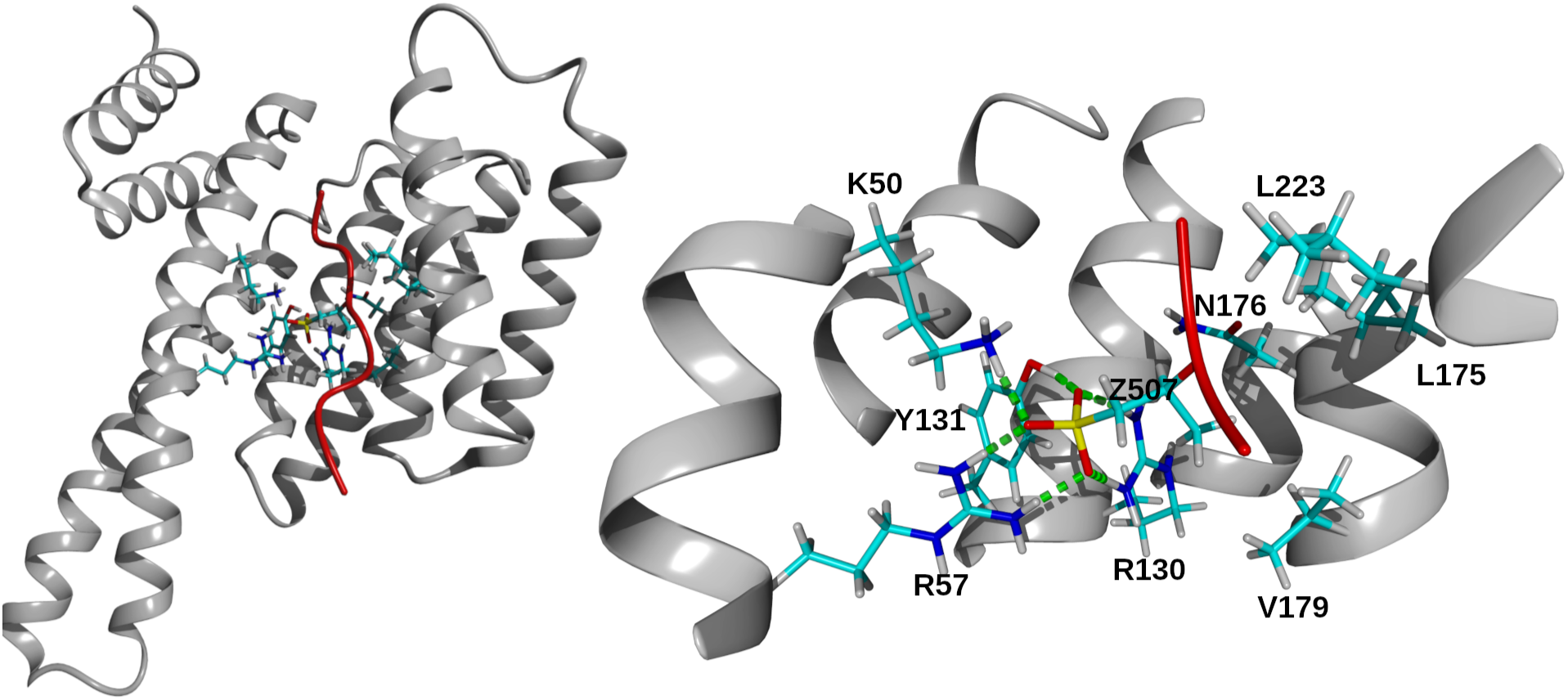
Residues of 14-3-3ε interacting with Pmb^507^. Left, structure of 14-3-3ε-[Pmb^507^]pT(502-510) complex. The secondary structure of protein and peptide is shown in gray ribbon and red tube, respectively. Interacting residues are in stick representation. Hydrogen, white; carbon, cyan; nitrogen, blue; oxygen, red and phosphor, yellow. Right, a close-up view showing residues of 14-3-3ε that form H-bonds with Pmb^507^ residue of [Pmb^507^]pT(502-510) peptide. Residues of 14-3-3ε are indicated with one letter code, and Pmb^507^ indicated by letter Z. H-bonds were identified within 0.4 nm distance between acceptor and donor atoms and indicated by green dashed line.

The binding of synthetic peptides to 14-3-3ε was initially assessed using DSF. The unfolding curves of 14-3-3 – peptide complexes and 14-3-3ε, and their corresponding first derivatives are shown in Figures 4 and S3. Analysis of DSF results indicated that [Pmb^507^]pT(502-510) peptide caused a ΔT_m_ of 14-3-3ε of 3.6°C, while all other peptides showed ΔT_m_ of 0.0°C. SPR was used to quantitatively determine K_D_ of the peptides to 14-3-3ε. Figures 5 and S4 show steady-state binding isotherm, and sensograms of the peptides binding to 14-3-3ε. The results showed that [Thr^507^](502-510) peptide did not bind to 14-3-3ε. [Glu^507^]pT(502-510), [sThr^507^]pT(502-510), and [Gla^507^]pT(502-510) had K_D_ of 141.75 ± 24.96 μM, 86.71 ± 8.07 μM, and 24.06 ± 8.43 μM, respectively. [Pmb^507^]pT(502-510) peptide had the highest affinity of 0.009 ± 0.001 μM (Table 1).

**Figure 4.**
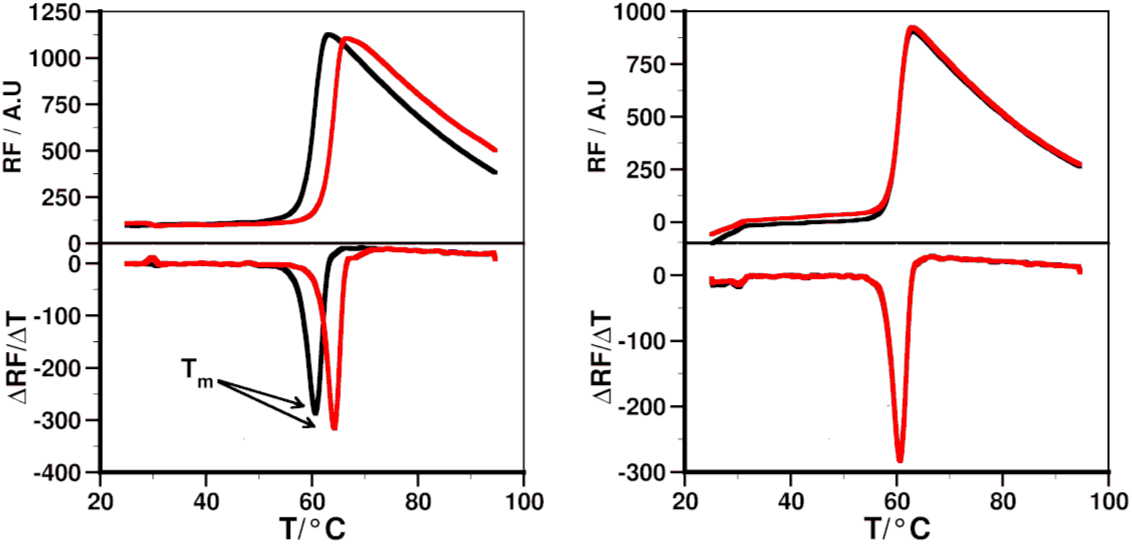
Thermal unfolding of 14-3-3ε protein, and its complexes with peptides. Thermal unfolding curve of 14-3-3ε without (black) and with peptide (red), and their respective first derivatives from differential scanning fluorimetry (DSF). Left panel, [Pmb^507^]pT(502-510); right panel, [sThr^507^]pT(502-510). The binding of the peptide is determined as the change in melting temperature (T_m_) between 14-3-3ε – peptide complex and 14-3-3ε alone.

**Figure 5.**
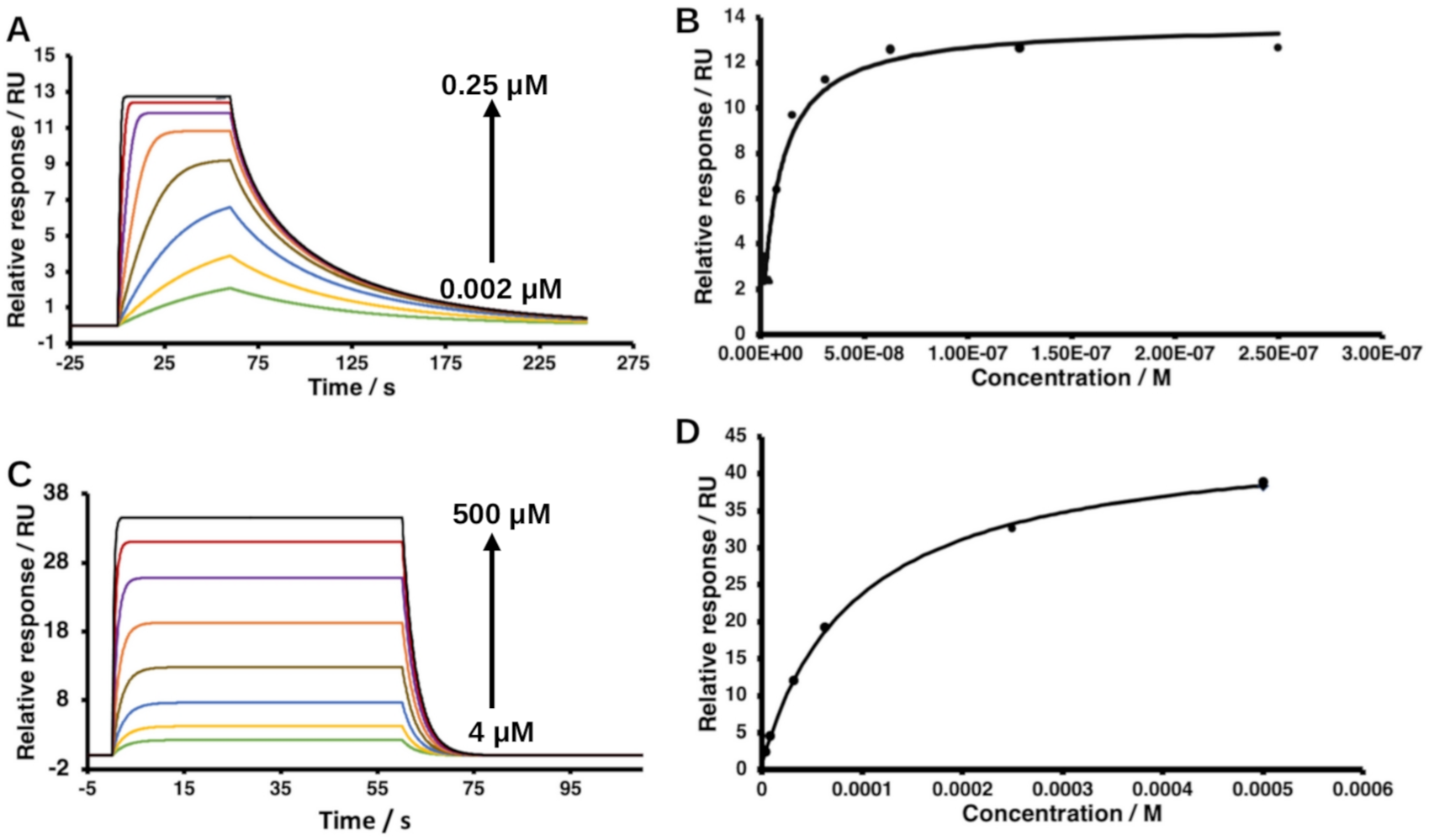
Surface plasmon resonance-based dose response binding of peptides to 14-3-3ε. A and B, sensograms and their corresponding binding isotherm of [Pmb^507^]pT(502-510); C and D, [sThr^507^]pT(502-510). The arrows indicate binding response of increasing concentrations of the peptides, concentrations range between 0.002 μM and 0.25 μM (green, 0.002 μM; yellow, 0.004 μM; blue, 0.008 μM; light brown, 0.016 μM; orange, 0.031 μM; purple, 0.063 μM; red, 0.125 μM; black, 0.25 μM) for [Pmb^507^]pT(502-510), and between 4μM and 500 μM (green, 3.9 μM; yellow, 7.8 μM; blue, 15.6 μM; light brown, 31.2 μM; orange, 62.5 μM; purple,125 μM; red, 250 μM; and black, 500 μM) for [sThr^507^]pT(502-510). 100 nM of 14-3-3ε was used in all experiments.

## Discussion

Interactions of 14-3-3 proteins with their binding partners have been primarily attributed to phosphorylated residues in the partners. However, 14-3-3 proteins have also been reported to bind proteins in phosphorylation-independent manner [24,27]. Therefore, here we studied the role of negative charges in binding partners for interaction with 14-3-3ε. We showed that phosphorylation is not necessary for binding of 14-3-3ε, but single or double negative charges are required. We used MD simulations to determine the stability of 14-3-3ε – peptide complexes.

All the peptides formed stable complexes with 14-3-3ε; no peptide dissociation was observed (Figure S1). The structural rearrangement was comparable to that of the phosphorylated peptides [30]. ISA calculations can be used to reveal interactions between binding partners [43] Our ISA results qualitatively confirmed that all the peptide had interactions with 14-3-3ε in the protein – peptide complexes. The Pmb residue of the highest affinity peptide, [Pmb^507^]pT(502-510), had the largest ISA, while the residues from non-binding and weakly binding peptides had smaller ISA.

Since the residues of 14-3-3 proteins that interact with the phosphorylated amino acid residues of their binding partners are known [32], we sought to determine if these interactions were maintained after residue substitutions in the parent peptide. In agreement with our previous results [19, 30], analysis of H-bonds involving pThr^507^ residue in the 14-3-3ε – pT(502-510) complex confirmed that the residues interacted with Arg^130^, Tyr^131^, Arg^57^ and Lys^50^ for the most part of the simulation time (Table S1). As expected, the uncharged residue Thr in [Thr^507^]pT(502-510) did not interact with the reported residues of the 14-3-3ε (Table S1). H-bond analysis further showed that the singly charged residues sThr^507^ and Glu^507^ in [sThr^507^] (502-510) and [Glu^507^]pT(502-510) peptides, respectively, exhibited short-lasting H-bonds with the residues, except the Arg^130^ residues that lasted for more than 85% of the simulation time. Gla^507^ of the [Gla^507^]pT(502-510) peptide interacted with Arg^130^ and Tyr^131^ for > 95% of the simulations time, but temporarily also interacted with Arg^57^ and Lys^50^. Like the pThr^507^, Pmb^507^ residue in [Pmb^507^]pT(502-510) peptide formed long-lasting interactions with all the four residues. The Pmb showed additional H-bonds with Asn^176^, Val^179^, Leu^175^ and Leu^223^. The H-bonds by residues in position 507 were formed by negatively charged side chain oxygen(s), this explains why no H-bonds were detected with Thr^507^ residue of pT(502-510) peptide, as well as the more and longer lasting interaction by the doubly charged residues than the singly charged residue. Pmb residues and pThr have larger VDW surface area (Figure S2), and subsequently larges ISAs, suggesting that besides charge, the larger of the side chain of residues in position 507 of the peptides might be of importance for the peptide binding to 14-3-3ε.

DSF studies showed that the [Pmb^507^]pT(502-510) peptide caused a shift of 3.6°C, while other peptides did not cause any shift in T_m_ of 14-3-3ε (Figure S3) at 30 μM. The concentration at which the parent peptide and its other analogs caused change in T_m_ of 14-3-3ε. The parent peptide, pT(502-510) shows ΔT_m_ of 3.7 °C [30]. Similar to DSF, in SPR experiments, the [Thr^507^]pT(502-510) did not bind 14-3-3ε at the studied concentrations (0–500 μM). This is in agreement with the fact that no interactions were observed between Thr^507^ residue of pT(502-510) peptide and residues of 14-3-3ε. This also indicates the crucial importance of negatively charged amino acid residues in peptide binders of 14-3-3ε. The [sThr^507^](502-510) and [Glu^507^]pT(502-510) had K_D_ of 86.71 ± 8.07 μM and 141.75 ± 24.96 μM, respectively. The peptide containing the doubly charged Gla, [Gla^507^]pT(502-510) showed a higher affinity (K_D_ of 24.06 ± 8.43 μM) (Table 1). Furthermore, the peptide with the doubly charged residue (Pmb) that has a phosphono group instead of phosphate group, [Pmb^507^]pT(502-510) showed the highest affinity (K_D_ of 0.009 ± 0.001 μM) for 14-3-3ε than all other analogs. [Pmb^507^]pT(502-510) also has higher affinity for 14-3-3ε than our previously reported best binder for 14-3-3ε [30]. The biophysical results are in agreement with our computational results showing stronger interactions between the [Pmb^507^]pT(502-510) peptide and 14-3-3ε. Similar to H-bond analysis, the [Gla^507^]pT(502-510) peptide showed weaker affinity than [Pmb^507^]pT(502-510) yet both Gla and Pmb are doubly charged, this could be due to different distributions of electrons on the side chain of these residues, resulting in different location of net charges, as well as the slight difference in their VDW surface areas.

Although binding of 14-3-3 with other proteins has been mostly reported with phosphorylated binding partners [20,21,44], it also interacts with unphosphorylated proteins like exoenzyme S [27,28], and unphosphorylated peptides [23,32]. In a study conducted by Petosa and colleagues [32] comparing binding of the unphosphorylated pan inhibitor of 14-3-3 proteins, R18 and the RAF-1 derived phosphopeptide pS-RAF-1-259, a crystal structure of 14-3-3ζ with R18 showed that a pentapeptide fragment of R18 (Trp-Leu-Asp-Leu-Glu) occupies binding site that overlaps with that of the phosphorylated peptides. The pentapeptide fragment contains two acidic amino acid residues (Asp and Glu), in the crystal structure, these residues are next to Lys^49^, Arg^56^, Arg^60^, and Arg^127^ residues of 14-3-3ζ, the phosphate group of pS-RAF-1-259 peptide is next to the same residues. In a different study on a peptide that contains two Asp residues [27], one Asp is located far (0.48 – 0.573 nm) from the basic residues of 14-3-3ζ to form significant interactions, the other Asp residue contacts Lys^49^. These studies are in agreement with our previous studies in which we reported similar residues (Lys^50^, Arg^57^, Arg^130^ and Tyr^131^) of 14-3-3ε to interact with the phosphorylated residue in our phosphopeptide analogs [19,31]. The same residues of 14-3-3ε interacted with the 507 residues in the peptides presented in this work.

In the current work, we demonstrated that negative charge(s), not necessarily phosphorylation, are required for binding to 14-3-3 proteins and that two negative charges are favored for high affinity binding. The isosteric replacement of pThr with sThr with one negative charge on the side chain resulted in weak binder peptide. We also designed a peptide, [Pmb^507^]pT(502-510), having two negatively charged side chain, that has the highest, nanomolar, affinity for 14-3-3ε. In our previous work we designed phosphopeptide analogs that inhibit squamous cell carcinoma cell growth [19]. Unlike these pervious analogs, the [Pmb^507^]pT(502-510) is likely more stable, since it is not susceptible to dephosphorylation, making it a better candidate for inhibition of CDC25A – 14-3-3ε interactions to promote apoptosis of cSCC cells.

## Author Contributions

S.L., and S.K., conceptualization; L.A.H. and S.L., funding acquisition; S.L., study design; S.K., computation, experiments and data analysis; S.K. and S.L., writing original draft, L.A.H., reviewing and editing.

## Acknowledgments

This work was supported by the National Institutes of Health R01 CA253573-01 and the state of Nebraska LB595 grants. Mass spectral analysis of synthetic peptides were performed by the Mass Spectrometry Core facility as a component of the Auditory Vestibular Technology Core within the Translational Hearing Center at Creighton University, School of Medicine funded by CoBRE Award 5P20GM139762 from the NIH.

