## Supporting Material for "Negative Charges, Not Necessary Phosphorylation, are Required for Ligand Recognition by 14-3-3 Proteins"

| | |
|---|---|
| *Corresponding author: | Sándor Lovas |
| Address: | Department of Biomedical Sciences |
| | Criss II, Room 313 |
| | Creighton University |
| | 2500 California Plaza |
| | Omaha, NE 68178 |
| Phone: | 402-280-5753 |
| Fax: | 402-280-2690 |
| E-mail: | |


**Running Title**: Charge Contribution to Recognition of 14-3-3 Proteins

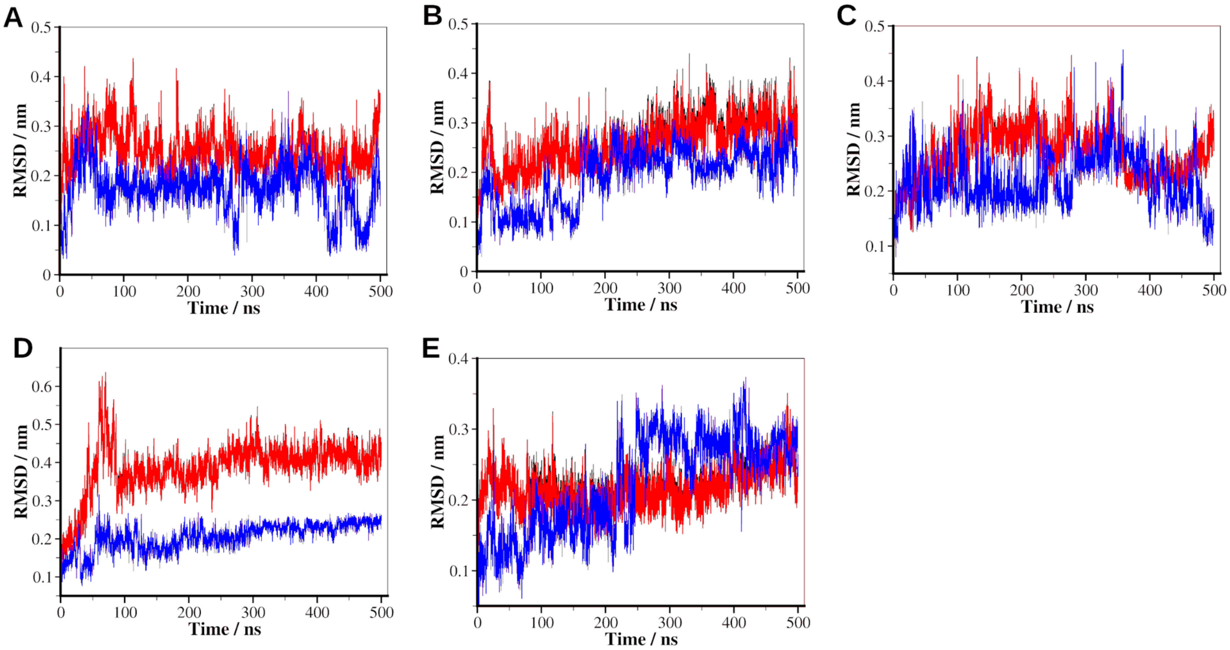


**Figure S1**. **Root mean square deviation (RMSD) analysis of simulations of 14-3-3ε – peptide complexes.** Time course RMSD of complexes during a 500 ns simulation; Black, 14-3-3ε-peptide complex; red, 14-3-3ε in the complex; blue, peptide in the complex. Since the RMSDs of complexes and the protein are overlapping, the black curve is hidden behind the red curve. (A) [Thr$^{507}$]pT(502-510); (B) [sThr$^{507}$](502-510); (C ) [Glu$^{507}$]pT(502-510); (D) [Gla$^{507}$]pT(502-510); (E) [Pmb$^{507}$]pT(502-510).

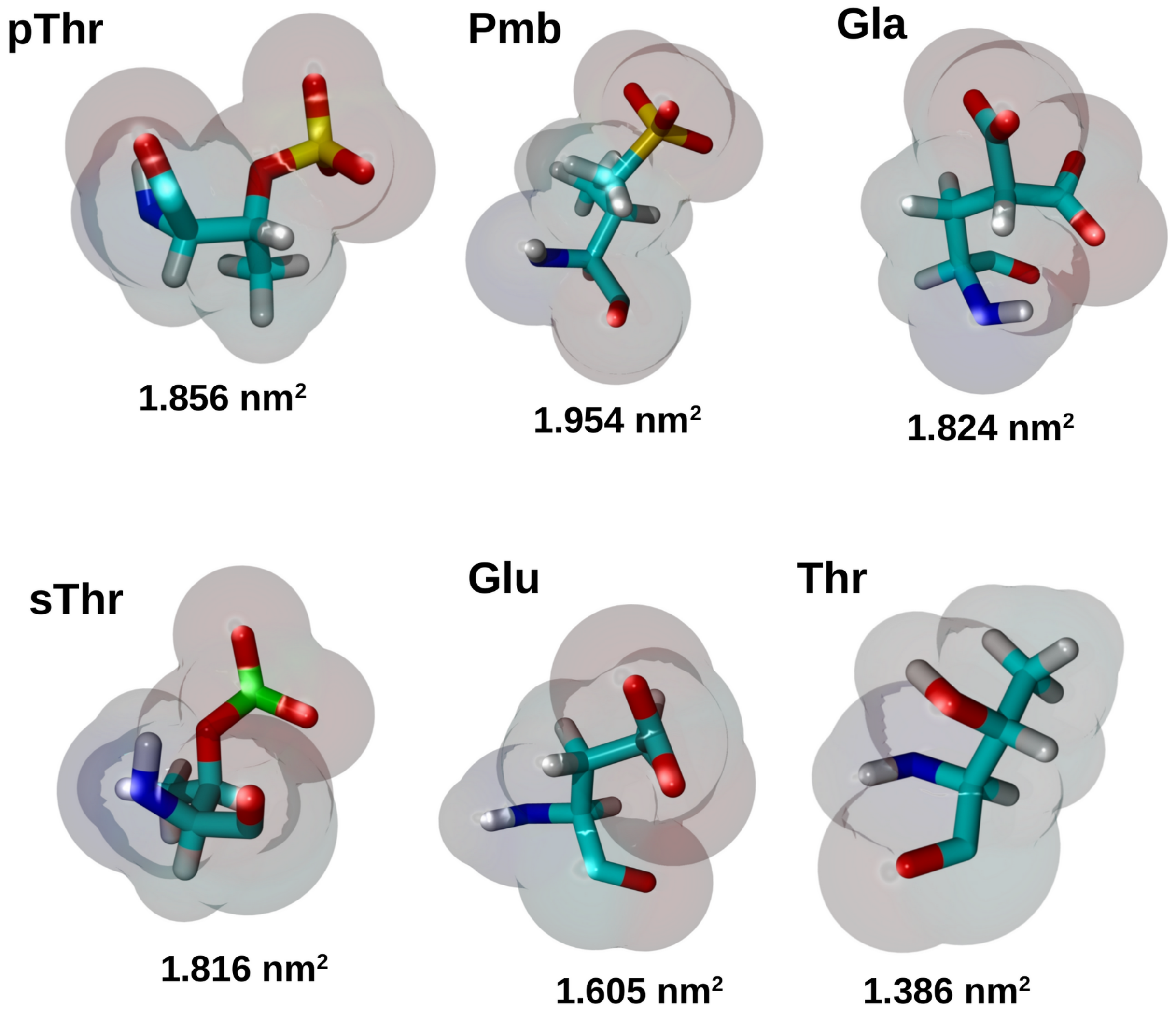


**Figure S2. Van der Waals (VDW) surface areas of amino acid residues**. The surface areas were calculated from representative structures from simulation trajectories.

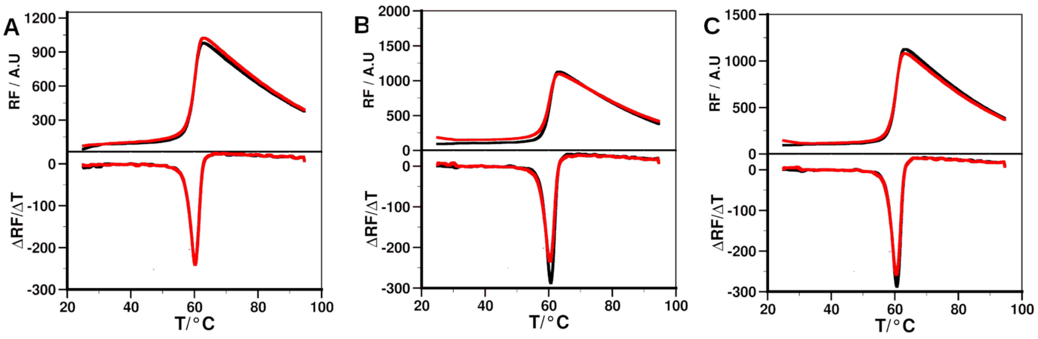


**Figure S3. Thermal unfolding of 14-3-3ε protein, and its complexes with peptides.** Melting curve of 14-3-3ε without (black) and with (red) peptide, and their respective first derivatives from differential scanning fluorimetry (DSF). Left panel, [Thr$^{507}$]pT(502-510); middle panel, [Glu$^{507}$]pT(502-510); right panel, [Gla$^{507}$]pT(502-510). Binding of the peptides is determined as the change in melting temperature (Tm) between 14-3-3ε – peptide complex and 14-3-3ε alone.

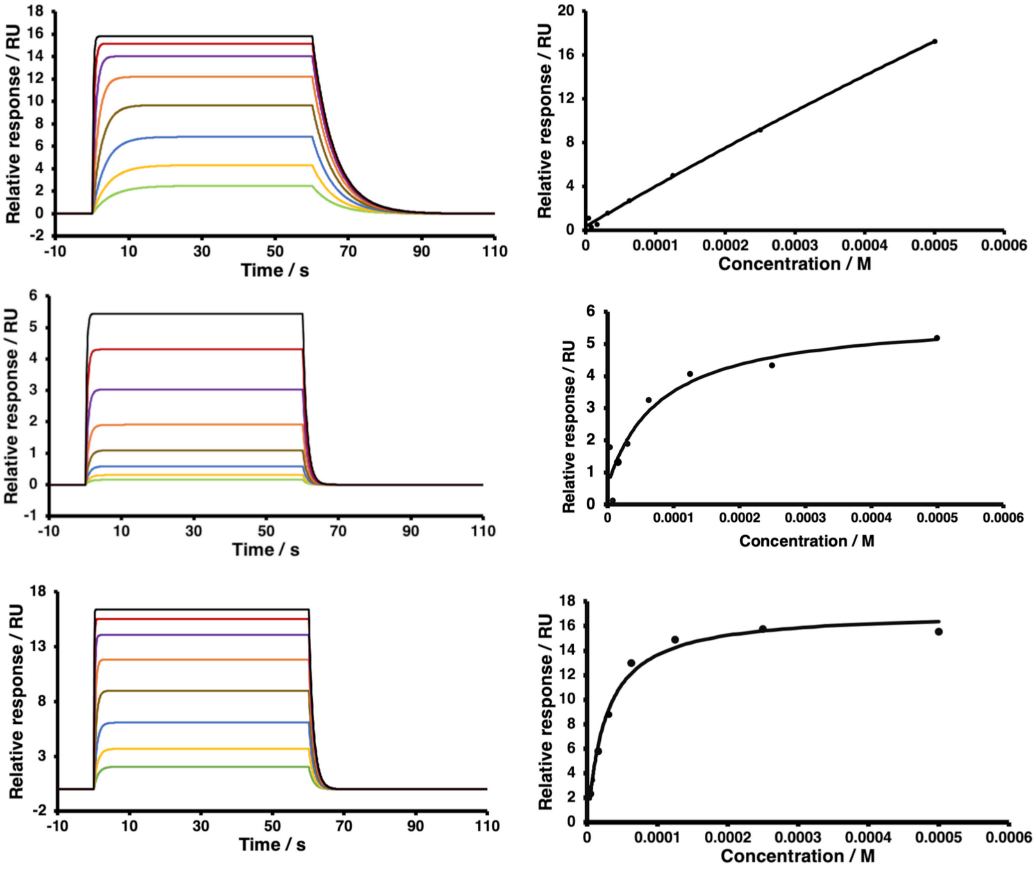


**Figure S4. Surface plasmon resonance-based dose response binding of the peptides to 14-3-3ε.** Sensograms and their corresponding binding isotherm. Upper panel, [Thr$^{507}$]pT(502-510); middle panel, [Glu$^{507}$]pT(502-510); lower panel, [Gla$^{507}$]pT(502-510). Peptide concentration range between 3.9 μM and 500 μM (green, 3.9 μM; yellow, 7.8 μM; blue, 15.6 μM; light brown, 31.2 μM; orange, 62.5 μM; purple,125 μM; red, 250 μM; and black, 500 μM), and 100 nM of 14-3-3ε was used in all experiments.

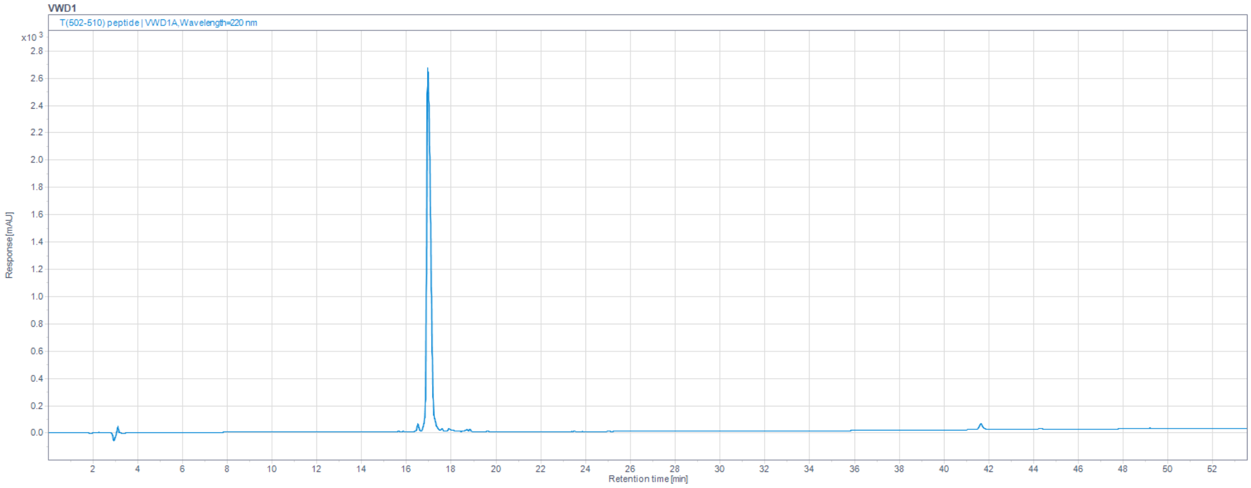


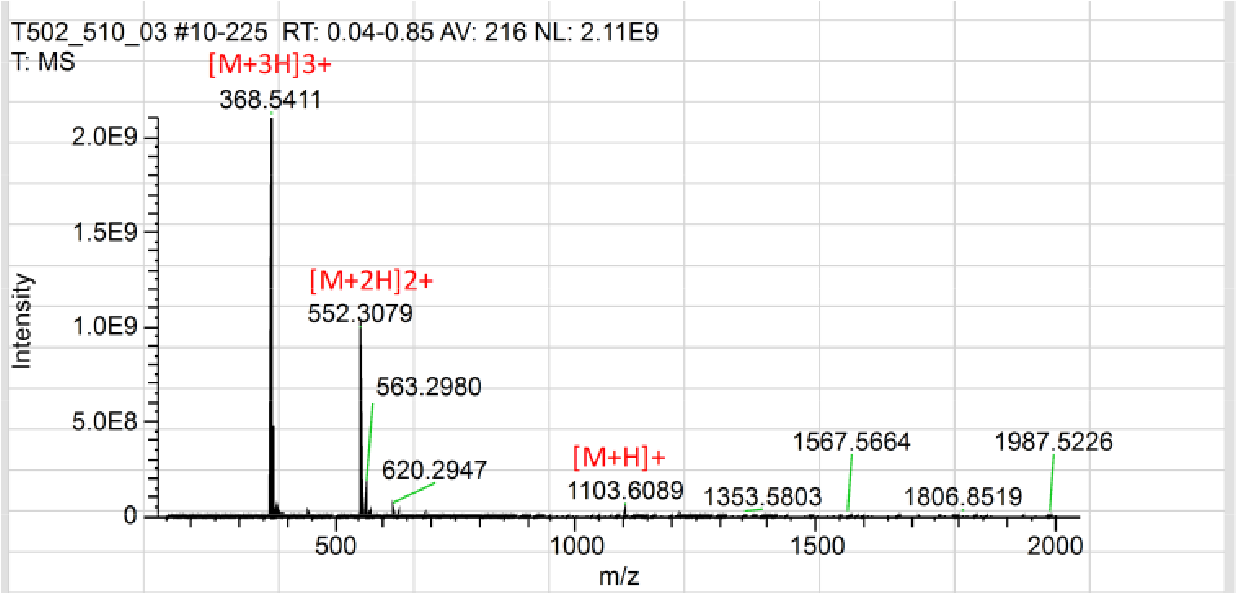


**Figure S5.** HPLC chromatogram and mass spectrum of the purified [Thr$^{507}$]pT(502-510).

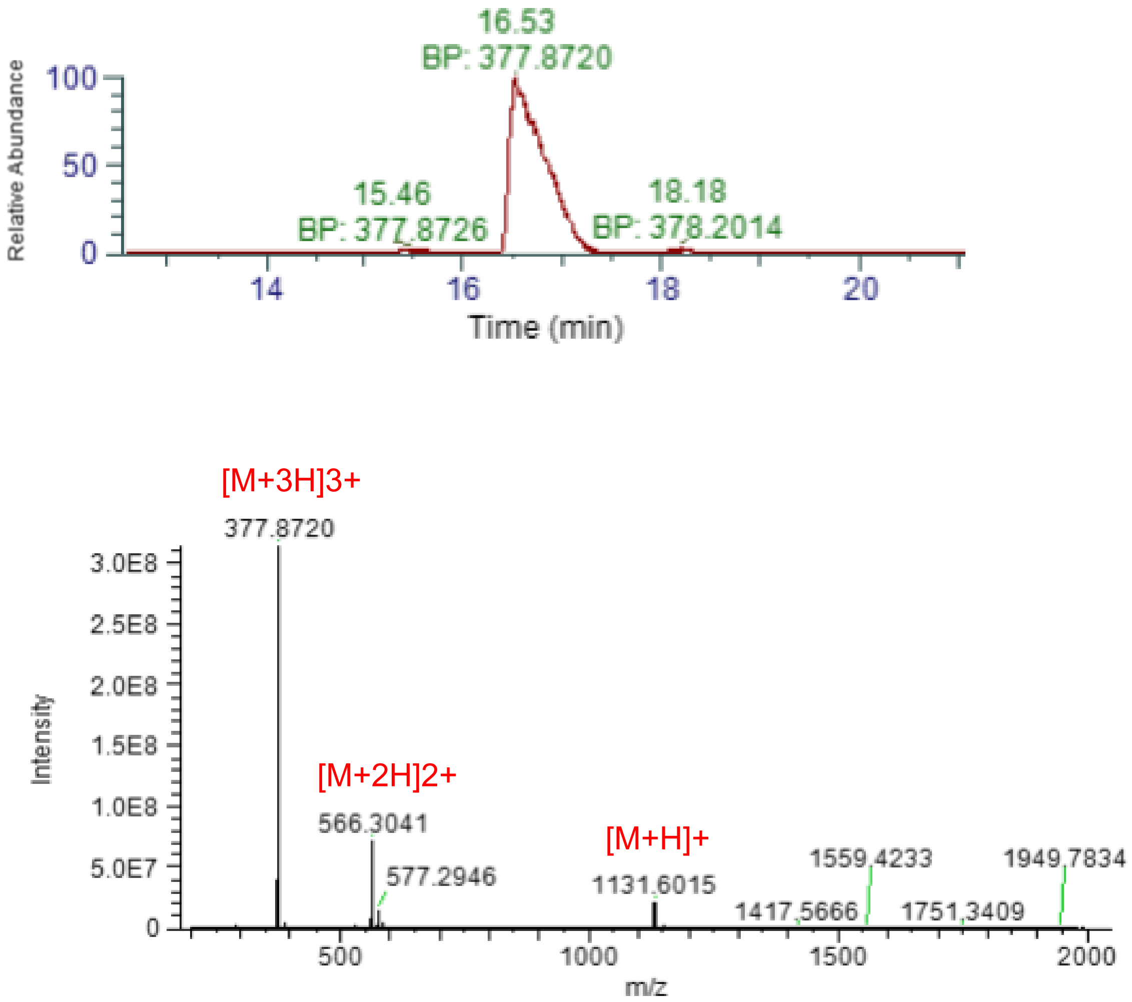


**Figure S6.** UHPLC chromatogram and mass spectrum of the purified [Glu$^{507}$]pT(502-510).

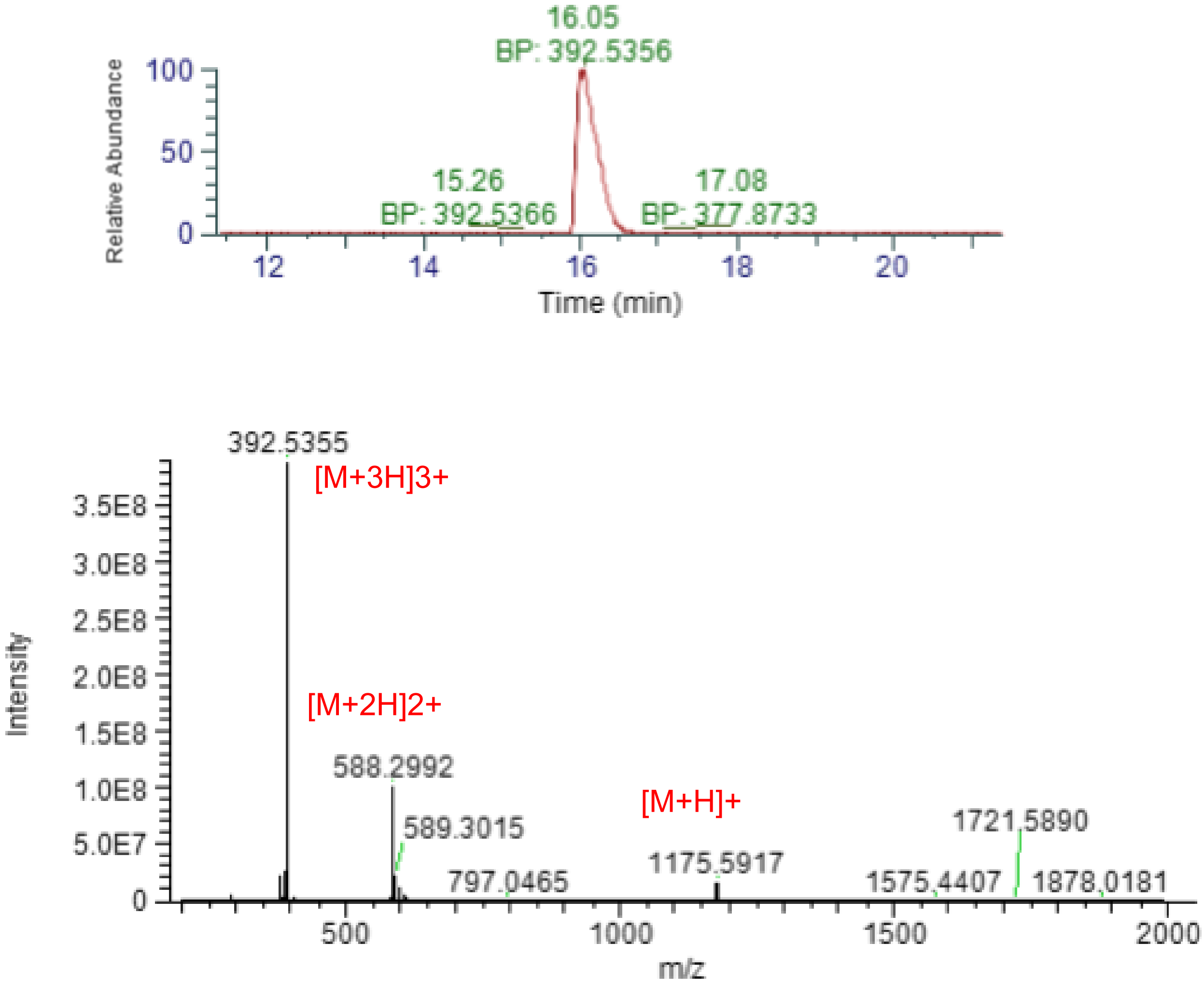


**Figure S7.** UHPLC chromatogram and mass spectrum of the purified [Gla$^{507}$]pT(502-510).

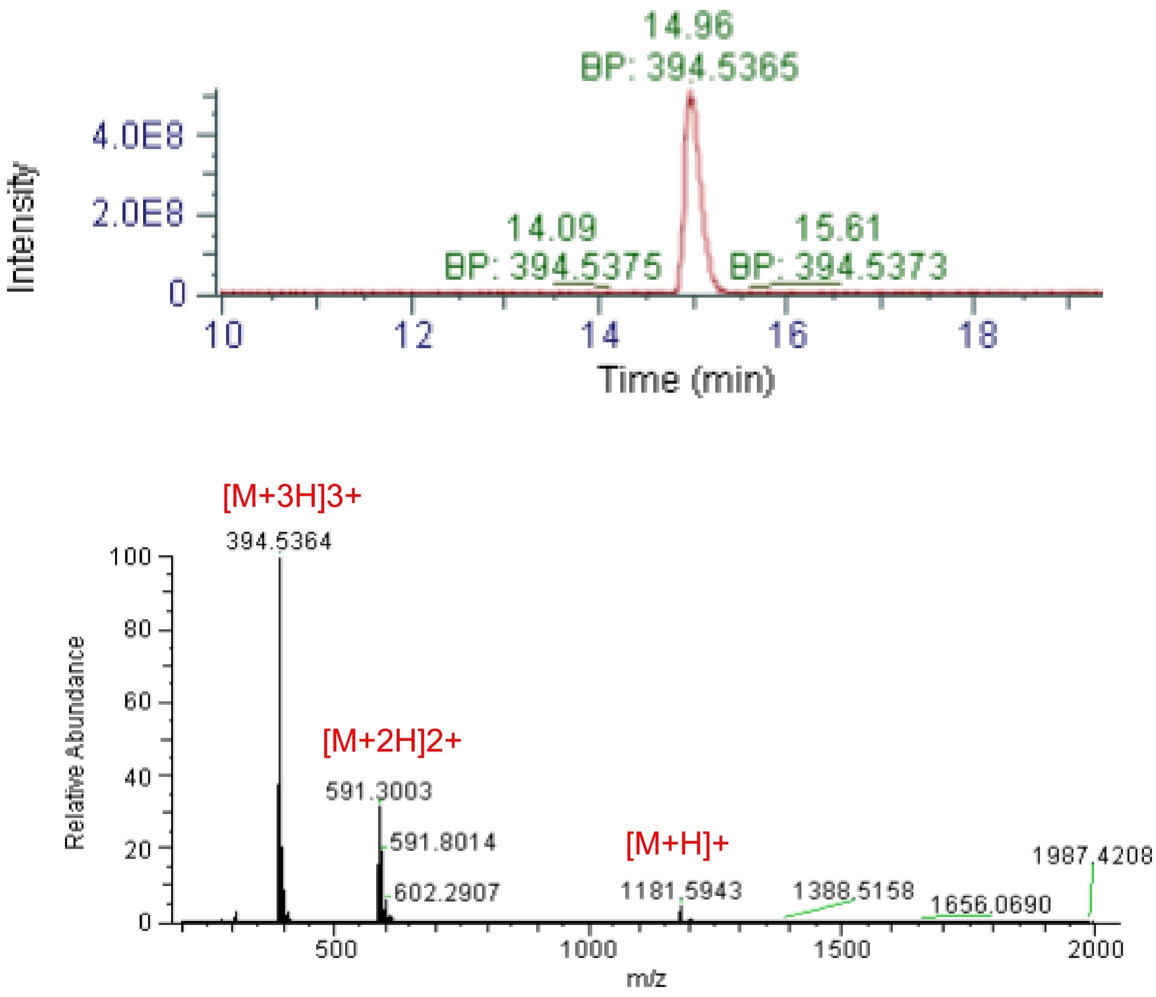


**Figure S8.** UHPLC chromatogram and mass spectrum of the purified [Pmb$^{507}$]pT(502-510).

**Table S1. H-bods between 507 residue of the peptides and 14-3-3ε.**

The bonds were calculated from 500 ns simulations. The donor-acceptor distance was 0.4 nm.

| Peptides | Residue of the protein | % occupancy |
|---|---|---|
| pT(502-510) | $Arg^{130}$ | 99.44 |
| | $Tyr^{131}$ | 98.36 |
| | $Arg^{57}$ | 97.94 |
| | $Lys^{50}$ | 43.85 |
| | $Ans^{176}$ | 0.22 |
| [$Thr^{507}$]pT(502-510) | No H-bonds detected | |
| [$sThr^{507}$]pT(502-510) | $Arg^{130}$ | 86.24 |
| | $Tyr^{131}$ | 9.13 |
| | $Lys^{50}$ | 1.06 |
| | $Arg^{57}$ | 0.80 |
| [$Glu^{507}$]pT(502-510) | $Arg^{130}$ | 87.78 |
| | $Tyr^{131}$ | 37.8 |
| | $Asn^{176}$ | 8.11 |
| | $Arg^{57}$ | 3.49 |
| | $Lys^{50}$ | 2.30 |
| [$Gla^{507}$]pT(502-510) | $Arg^{130}$ | 98.36 |
| | $Tyr^{131}$ | 96.41 |
| | $Lys^{50}$ | 23.28 |
| | $Arg^{57}$ | 3.97 |
| | $Arg^{61}$ | 0.78 |
| [$Pmb^{507}$]pT(502-510) | $Arg^{130}$ | 100 |
| | $Arg^{57}$ | 99.9 |
| | $Tyr^{131}$ | 99.16 |
| | $Lys^{50}$ | 65.54 |
| | $Asn^{176}$ | 19.33 |
| | $Val^{179}$ | 12.02 |
| | $Leu^{223}$ | 2.96 |
| | $Leu^{175}$ | 2.02 |

**Table S2. Masses of the peptide analogs as determined by mass spectrometry.**

| Peptide | Expected mass ($[M+H]^+$) m/z | Observed mass ($[M+H]^+$) m/z |
|---|---|---|
| [Thr$^{507}$]pT(502-510) | 1103.5963 | 1103.6089 |
| [Glu$^{507}$]pT(502-510) | 1131.6018 | 1131.6015 |
| [Gla$^{507}$]pT(502-510) | 1175.5916 | 1175.5917 |
| [Pmb$^{507}$]pT(502-510) | 1181.5939 | 1181.5943 |
